# Endurance exercise drives temporal and sexual dimorphic multi-omic adaptations in liver metabolism-Findings from MoTrPAC

**DOI:** 10.1101/2025.05.16.652859

**Authors:** Taylor J. Kelty, Edziu Franczak, Nicole R. Gay, Gina M. Many, Tyler J. Sagendorf, James A. Sanford, Zhenxin Hou, David A. Gaul, Facundo M. Fernández, Charles Burant, Andrea L. Hevener, Joshua N. Adkins, Sue Bodine, Malene E. Lindholm, Eric A. Ortlund, Simon Schenk, John P. Thyfault, R. Scott Rector, the MoTrPAC Research Group

## Abstract

The mechanisms by which exercise modulate liver metabolism, a central regulator of systemic metabolism, are poorly understood. Leveraging data from MoTrPAC, we analyzed liver adaptations across 1, 2, 4, and 8 weeks of exercise in male and female rats using multi-omic approaches. Female livers displayed a progressive increase in oxidative phosphorylation (OXPHOS) complexes (at the protein level), while male livers showed an increase in acetylation of OXPHOS, TCA cycle, and fatty acid oxidation enzymes. Exercise also enhanced liver cholesterol and bile acid synthesis, reducing liver lipid metabolites in males after 8 weeks of exercise. Male rats had higher fecal cholesterol and cholic acid levels, indicating a sex-specific mechanism of lipid excretion with exercise. Moreover, 8 weeks of training reduced markers related to hepatic stellate cell activation and fibrosis in both sexes. This study highlights the sexual dimorphic and temporal molecular signatures by which exercise modulates liver metabolism to provide hepatoprotective effects.

---

Cardiorespiratory fitness is the most significant predictor for all-cause mortality in both men and women^1,2,21,2^. Endurance exercise has been shown to improve parameters of metabolic health^3^, particularly in patients with metabolic syndromes^4,5^, serving as an accessible frontline defense against metabolic diseases, including metabolic-dysfunction-associated steatotic liver disease (MASLD)^6–8^ colloquially known as fatty liver disease. During periods of metabolic stress (i.e., exercise, fasting), the liver modulates its uptake (fatty acids, lactate, amino acids, etc.) and secretion (glucose and ketones) of metabolites in response to metabolic and endocrine signaling to meet systemic energetic needs^9,10^. Importantly, exercise training imparts a durable liver adaptation, promoting tighter coupling of ATP synthesis to glucose production at any given workload^11–13^. Because the liver is critical in orchestrating substrate availability to tissues during exercise and is central to metabolic regulation, understanding exercise-induced molecular adaptations in the liver is critically important^14–16^.

Rodent studies from the last half century have revealed a myriad of exercise-induced liver adaptations ranging from markers of elevated mitochondrial function^12,17,18^ to robust changes in glucose metabolism^10,19–21^. Further research has corroborated and enhanced these findings to reveal that exercise can effectively prevent and reverse MASLD in both preclinical models and humans^22–28^. Omic-based approaches have recently been employed to better understand exercise-induced liver adaptations in transcripts^25^, proteins^26,29–31^, and metabolites^27,28^. Moreover, a substantial proportion of studies have focused on the male sex, despite evidence of a sexual dimorphism in liver mitochondrial dynamics^11,13,32–34^. Here, this study applied multi-omic analyses to capture temporal liver adaptations to exercise.

The purpose of the Molecular Transducers of Physical Activity Consortium (MoTrPAC)^35^ is providing detailed maps of the multi-omic response to acute physical activity and exercise training across tissues^36^. We have recently published findings on the multi-omic changes across tissues in exercise-trained (1, 2, 4, or 8 weeks of treadmill running) female and male rats (aged 6 months)^35,37^. In this cohort of rats, we characterized the temporal training response related to mitochondrial analytes^38^, transcriptomic and epigenetic signatures^39^, complex trait genetics^40^, and adiposity^41^. Utilizing the data generated by MoTrPAC^35^, we herein applied an integrative multi-omic methodology to focus on the characterization of the liver exercise training response in both male and female rats at 1, 2, 4, and 8 weeks of training. These findings provide valuable insight into how chronic exercise impacts liver metabolism, outcomes that can be leveraged to develop treatments that targets optimal liver health and prevent or treat MASLD.

## RESULTS

### Sex-specific and temporal liver remodeling is associated with enhanced mitochondrial and lipid metabolism following chronic exercise training

To identify the most prominent temporal or sex-associated liver exercise adaptations, we first examined each ome (transcripts, proteins, metabolites, lipids, and some molecular modifications) in response to progressive exercise training (Fig. 1A). In this graphical cluster, edges are split by each of the top 4 impacted omes analyzed (unable to identify a transcript trajectory due to relatively limited transcript adaptations across time), where each node represents 1 of 9 states (row labels) at each of the 4 sampled training timepoints (column labels). Edges are drawn between adjacent nodes to represent the path of differential analytes over the training time course. This graph includes the largest paths with both node and edge size being proportional to the number of analytes represented. Of the top liver clustering trajectories from each ome, endurance training induced a robust increase in clustering in the acetylome over the first 4 weeks of exercise training in males (increased at eight weeks in both males and females), decreased many proteomic clusters after 8 weeks of exercise in both males and females, and a time- and sex-specific downregulated clustering of phosphoproteome (males decreased in the first 2 weeks, both at 4 weeks, and females decreasing at training week 8), and decreased metabolomic analytes at 8 weeks in males.

**Figure 1:**
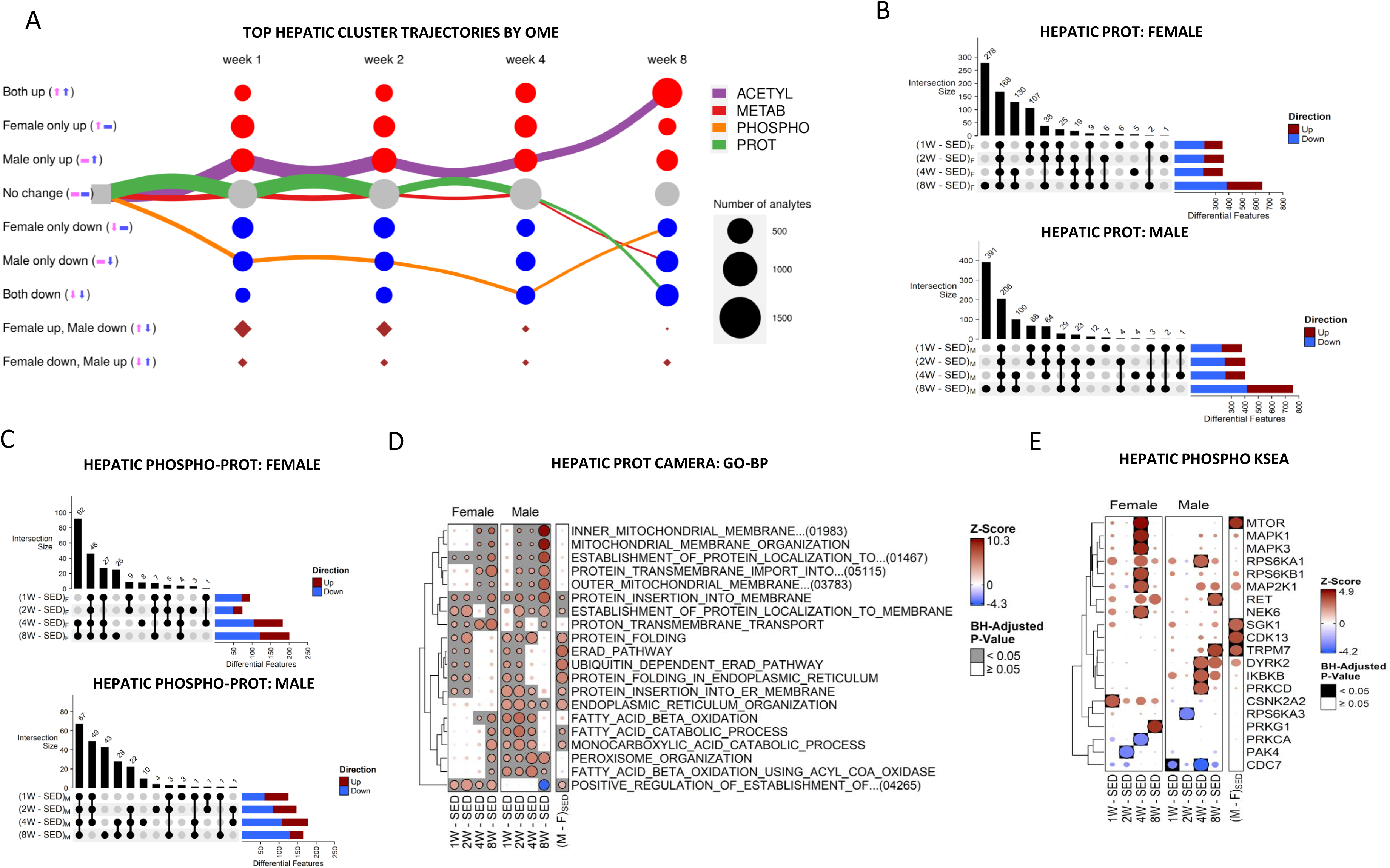
Sex-specific and temporal liver remodeling is associated with enhanced mitochondrial and lipid metabolism following chronic exercise training. **A)** Graphical representation of training-differential analytes in liver tissue. This graph includes the largest paths with both node and edge size being proportional to the number of analytes represented. **B-C)** Upset plots of statistically significant (FDR < 0.05) proteins **(B)** and phosphoproteins **(C)** between age-matched trained and sedentary females and males. **D)** Bubble heatmaps depicting top biological processes (BP) from the gene ontology (GO) database across training and sedentary conditions. Heatmaps were derived from the proteomic CAMERA-PR results. Circles are colored according to the set-level z-score and scaled by row so that the most significant comparison is of maximum area. **E)** Inferred activity of the indicated kinases across eight weeks of training in liver tissue. Blue bubbles indicate a decrease and red indicated increase in endurance training compared to sedentary rats or a decrease in sedentary male compared to sedentary female rats. A gray or black background indicates a significant result (adjusted p-value < 0.05).

Exercise training induced distinct temporal patterns between differentially regulated proteins, representing a combined 794 (female) and 914 (male) differentially abundant proteins (DAPs) and 227 (female) and 233 (male) phosphosites across all time points in false discovery rate (FDR) < 0.05, Fig. 1B-C). Notably, 278 (female) and 391 (male) of these DAPs and 25 (female) and 43 (male) phosphosites were unique to 8 weeks of exercise training suggestive of a cumulative effect of training on the proteome and phosphoproteome. Additionally, 168 (female) and 206 (male) DAPs and 92 (female) and 67 (male) phosphosites were observed across all exercise timepoints (1,2,4, 8 weeks). To bridge our understanding between differentially expressed analytes and related biological functions, we performed Pre-Ranked Correlation Adjusted MEan RAnk gene set testing (CAMERA-PR)—a method that accounts for inter-gene correlation to more correctly control the false positive rate—to test for enrichment of Gene Ontology (GO) Biological Processes (BP)^47–50,52^ (Fig. 1D and E). Both female and male liver proteomes were positively enriched for terms associated with mitochondrial metabolic pathways with chronic exercise training (mitochondrial membrane organization and fatty-acid beta-oxidation), with the male liver acquiring these adaptations earlier (week 2 of training) compared to females who acquired them later (week 4 of training) (Fig. 1D). The slower adaptation in females is likely attributable to females displaying enhanced mitochondrial oxidative phosphorylation (OXPHOS) profiles in the sedentary condition compared to males. CAMERA-PR was also applied to sets of phosphosites grouped according to their known kinases to examine changes in predicted kinase activity. We refer to the use of CAMERA-PR in conjunction with kinase sets as Kinase–Substrate Enrichment Analysis (KSEA). The female liver phosphoproteome displayed a robust increase in mammalian target of rapamycin (mTOR) after 4 weeks of training (Fig. 1E). mTOR1 activates *de novo* lipogenesis through downstream activation of sterol regulatory element binding protein 1c (SREBP1c)^63,64^ corroborated by transcriptomic GO-BP analysis. The male liver phosphoproteome showed a robust increase in MAP kinase signaling at 4 weeks, while the female liver phosphoproteome displayed a decrease in Cell division cycle 7 (CDC7) signaling at 4 weeks (Fig. 1E). The proliferative effects of Mitogen-activated protein (MAP) kinase signaling in the liver is well documented^65,66^, while CDC7 inhibition is reported to have anti-fibrotic effects^67^, consistent with Ingenuity Pathway Analysis (IPA) predictions of decreased stellate cell proliferation and fibrosis from 8 weeks of training in Fig.2.

**Figure 2:**
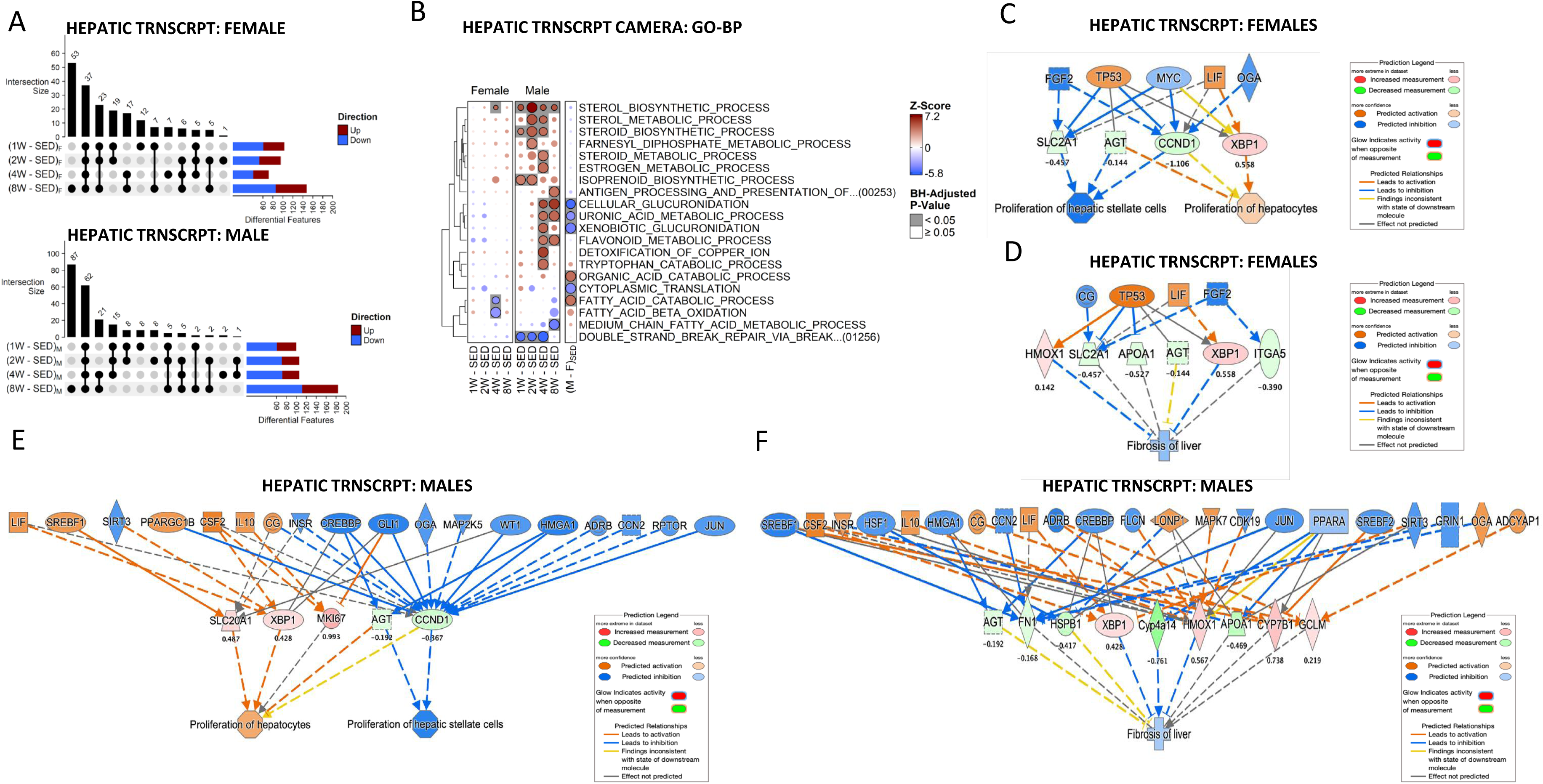
Chronic exercise altered liver transcriptome to promote sterol metabolism in hepatocytes, inhibiting hepatic stellate cell activation independent of sex. **A)** Upset plots of statistically significant (FDR < 0.05) transcripts between age-matched trained and sedentary females and males. **B)** Bubble heatmap depicting top biological processes (BP) from the gene ontology (GO) database across training and sedentary conditions. The heatmap was derived from the transcriptomic datasets using the CAMERA-PR method. Blue bubbles indicate a decrease and red an increase in endurance training compared to sedentary rats or a decrease in sedentary male compared to sedentary female rats. A gray or black background indicates a significant result (adjusted p-value < 0.05). **C and D)** Predicted decreased proliferation of stellate cells and increased proliferation of hepatocytes **(C)** and decrease in fibrosis **(D)** of the liver after 8 weeks of treadmill running in females. **E and F)** Decreased proliferation of stellate cells and increased proliferation of hepatocytes **(E)** and decrease in fibrosis **(F)** of the liver after 8 weeks of treadmill running in males. DEGs take all timepoints and sexes into account (FDR<0.05). Blue and green colors represent a decrease and orange, and red colors represent an increase in endurance training compared to sedentary rats (n=5). Expression levels of DEGs are shown below each node and displayed as log_2_FC (fold change). DEGs with z-score ≥ 2 were considered activated.

### Chronic exercise altered liver transcriptome to promote sterol metabolism in hepatocytes, inhibiting hepatic stellate cell activation independent of sex

Exercise training-induced distinct temporal patterns between differentially regulated transcripts, representing a combined 192 and 226 downregulated differentially expressed genes (DEGs) across all time points in females and males, respectively (FDR < 0.05, Fig. 2A). Notably, 53 (female) and 87 (male) of these DEGs were unique to 8 weeks of exercise training. Additionally, 37 (female) and 62 (male) DEGs were observed across all exercise timepoints (1,2,4, 8 weeks). We utilized CAMERA-PR to test for enrichment of Gene Ontology (GO) Biological Processes (BP) on the transcriptome. Across 8 weeks of exercise training, the male liver transcriptome was positively enriched for terms related to sterol biosynthetic and metabolic processes (Fig. 2B). Sterol biosynthetic processes were also enriched for the females but was not observed until after the 4^th^ week of exercise training (Fig. 2B). IPA was used to generate mechanistic networks based on top-predicted biological functions and upstream regulators following 1, 2, 4, or 8 weeks of exercise training. Differentially expressed analytes (i.e genes or proteins) and corresponding log_2_ fold changes were used as IPA input. IPA compared uploaded genes and proteins to a curated knowledge base of molecular relationships derived from peer-reviewed literature and public databases to identify enriched pathways and upstream regulators. IPA predicted decreased hepatic stellate cell (HSC) activation and fibrosis with increased hepatocyte proliferation, independent of sex (Fig. 2C-F). DEGs identified related to HSC activation and fibrosis in both males and females included angiotensinogen (AGT) and cyclin D1 (CCND1). Males showed a downregulation of AGT (log_2_FC = −0.192) and CCND1 (log_2_FC = −0.367) and females showed a downregulation AGT (log_2_FC = −1.106) and CCND1 (log^2^FC = −0.144). AGT and CCND1 is converted to angiotensin in the liver^95^, which promotes HSC activation and fibrosis through profibrotic pathways such as transforming growth factor beta-1 (TGF-β1) signaling and oxidative stress^96,97^. CCND1 regulates cycle progression from G1 to S phase and is associated with proliferation and activation of HSCs^99^.

### Female-specific liver proteome upregulation of OXPHOS complexes, carbohydrate metabolism, and amino acid metabolism with chronic exercise training

Based on the proteomics enrichment analysis results, we focused on baseline differences in mitochondria as well as mitochondrial adaptations to chronic exercise. Filtering transcriptomic and proteomic analytes using MitoCarta3.0 revealed significant upregulation of metabolic processes associated with liver mitochondria, that was limited at the transcript level (Fig. 3A), but robust at the protein level (Fig. 3B). While total mitochondrial protein content was similar between sedentary male and female rats (Extended Data Fig. 1A), significant divergence in abundance appeared between male and female sedentary rats. Sedentary male rats showed greater average abundance of mitochondrial proteins involved in fatty acid oxidation, carbohydrate metabolism, ketone metabolism, and branched-chain amino acid metabolism (p<0.05, Fig. 3B) compared to sedentary females. Overall total abundance of OXPHOS proteins was similar between sedentary males and females; however, complex I was significantly higher in sedentary female rats compared to males (p<0.05).

**Figure 3:**
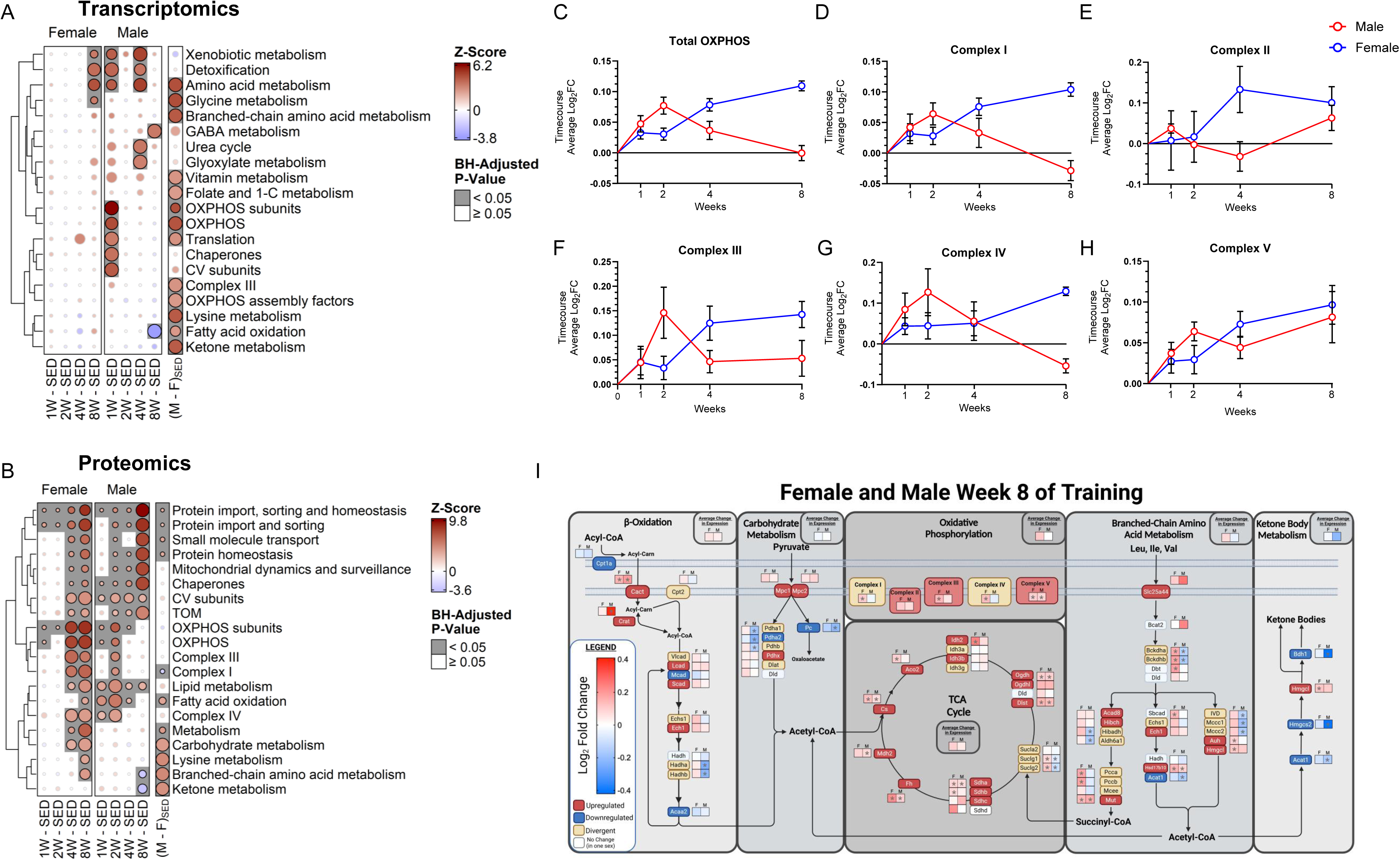
Female-specific liver proteome increases in abundance of OXPHOS complexes, carbohydrate metabolism, and amino acid metabolism with chronic exercise training. Bubble heatmaps indicating the direction and magnitude of change in abundance of the top 20 most significantly altered pathways within the **A)** transcriptome and **B)** proteomic differences following exercise training and at sedentary levels. **C)** Time course training effect on average log_2_FC protein abundance of all proteins involved in OXPHOS. **D-H)** Change in average protein log_2_FC abundance during the 1, 2, 4, and 8 weeks of training for each complex of OXPHOS. **I)** Log_2_FC in protein abundance at 8 weeks of training for all significantly altered metabolic process within liver mitochondria generated via BioRender. Protein labels in red represent increases in abundance and blue represent decreases in abundance within both male and female rats. Yellow labeled proteins show divergent change in direction of abundance between males and females. White labels represent no change in abundance within at least one sex (log_2_FC <0.01). Significant protein abundance change compared to sedentary conditions within sex is presented by *, p<0.05. Values are represented as mean ± standard error.

Exercise training promoted a sustained, progressive increase in mitochondrial protein abundance in female livers, including proteins involved in OXPHOS, fatty acid oxidation, branched-chain amino acid catabolism and carbohydrate metabolism (p<0.05, Fig. 3B). Inversely, exercise training failed to promote significant changes in the abundance of liver mitochondrial proteins involved within these same pathways in males, other than fatty acid oxidation and OXPHOS proteins showing significantly increased protein abundance during the initial 2 weeks of training (p<0.05) before falling back to near sedentary levels for the remainder of the training period (Fig. 3B). Interestingly, exercise promoted a significant reduction in proteins involved in ketone body synthesis and branched-chain amino acid catabolism following the 8-week training period. Both sexes showed a robust increase in the abundance of proteins involved in protein import into the mitochondria during all training phases (p<0.05). Sedentary transcript levels were congruent with protein abundance, yet the exercise-mediated alteration in the mitochondrial proteome was not reflected at the transcript level (Fig. 3A). Livers were collected 2 days after the last bout of endurance exercise training, thus the dampened transcript response is likely attributed to the livers rapid turnover of mRNA compared to other metabolically active tissues^68^.

Male and female rats displayed differential responses to exercise for proteins that compose the OXPHOS system (Fig. 3C). In females, protein abundance of many of the OXPHOS complexes (Fig. 3D-H) increased throughout the duration of the training period, resulting in an overall average increase in OXPHOS protein abundance of 7.8% (average log_2_FC = 0.11, Fig. 3C). In stark contrast, males displayed divergent training responses for OXPHOS protein abundance with exercise training. While complexes I, III, IV, and V displayed significantly increased abundance during the initial 2 weeks of exercise training (p<0.05), complex V was the only complex that showed a sustained elevation in abundance from sedentary levels throughout the entire 8-week training period in males (Fig. 3D-H).

Cumulatively, the female liver proteome appears to respond more robustly to chronic exercise training than males, driving greater protein abundance changes in metabolic (lipid, carbohydrate, branched-chain amino acids) pathways and OXPHOS content within the mitochondria (Fig. 3I). While most proteins within these pathways show similar directional change in abundance in response to exercise (depicted as blue-down regulated, red-upregulated), several proteins show sexual divergence in exercise-mediated abundance (depicted as yellow). This includes abundance of hydroxyacyl-CoA dehydrogenase trifunctional multienzyme complex subunit alpha (Hadha) and hydroxyacyl-CoA dehydrogenase trifunctional multienzyme complex subunit beta (Hadhb) within fatty acid beta-oxidation, Pyruvate Dehydrogenase E1 Subunit Alpha 1 (Pdha1) and pyruvate dehydrogenase E1 subunit beta (Pdhb) in carbohydrate metabolism, succinate-CoA ligase GDP/ADP-forming subunit alpha (Sulcg1) and succinate-CoA ligase GDP/ADP-forming subunit beta (Sulcg2) within the TCA cycle, Branched Chain Keto Acid Dehydrogenase E1 Subunit alpha (Bckdha) and Branched Chain Keto Acid Dehydrogenase E1 Subunit Beta (Bckhdb) in branched-chain amino acid catabolism, and complexes I and IV within OXPHOS. For each of these proteins, females had higher abundance while males significantly decreased abundance. Thus, while exercise impacts the abundance of various proteins uniformly, several show sex-dependent responses to exercise training.

### Sex-specific increase in OXPHOS, TCA cycle, and fatty acid oxidation enzymes in male liver acetylome with chronic exercise training

Our previous report demonstrated that exercise increased mitochondrial protein acetylation in the liver, particularly affecting proteins involved in OXPHOS, TCA cycle, fatty acid oxidation, and branched-chain amino acid catabolism^38^. Here, we extended the earlier analysis by examining innate sex differences and the progressive impact of exercise training on liver mitochondrial acetylation. Despite there being limited differences in overall acetylation levels under sedentary conditions between sexes (Extended Data Fig. 1A), protein-specific enrichment for acetylation correlated with sex differences in total protein abundance. In female sedentary rats, there was a 13% greater accumulation of acetylation events on OXPHOS proteins (log_2_FC = 0.18, Fig. 4A), whereas males exhibited higher levels of acetylated proteins in the TCA cycle (11% greater, log_2_FC = 0.15) and branched-chain amino acid catabolism pathways (17% greater, log_2_FC = 0.23; Fig. 4A). Sedentary female rats demonstrated greater overall acetylation levels across all complexes within OXPHOS, excluding complex II (Fig. 4B).

**Figure 4:**
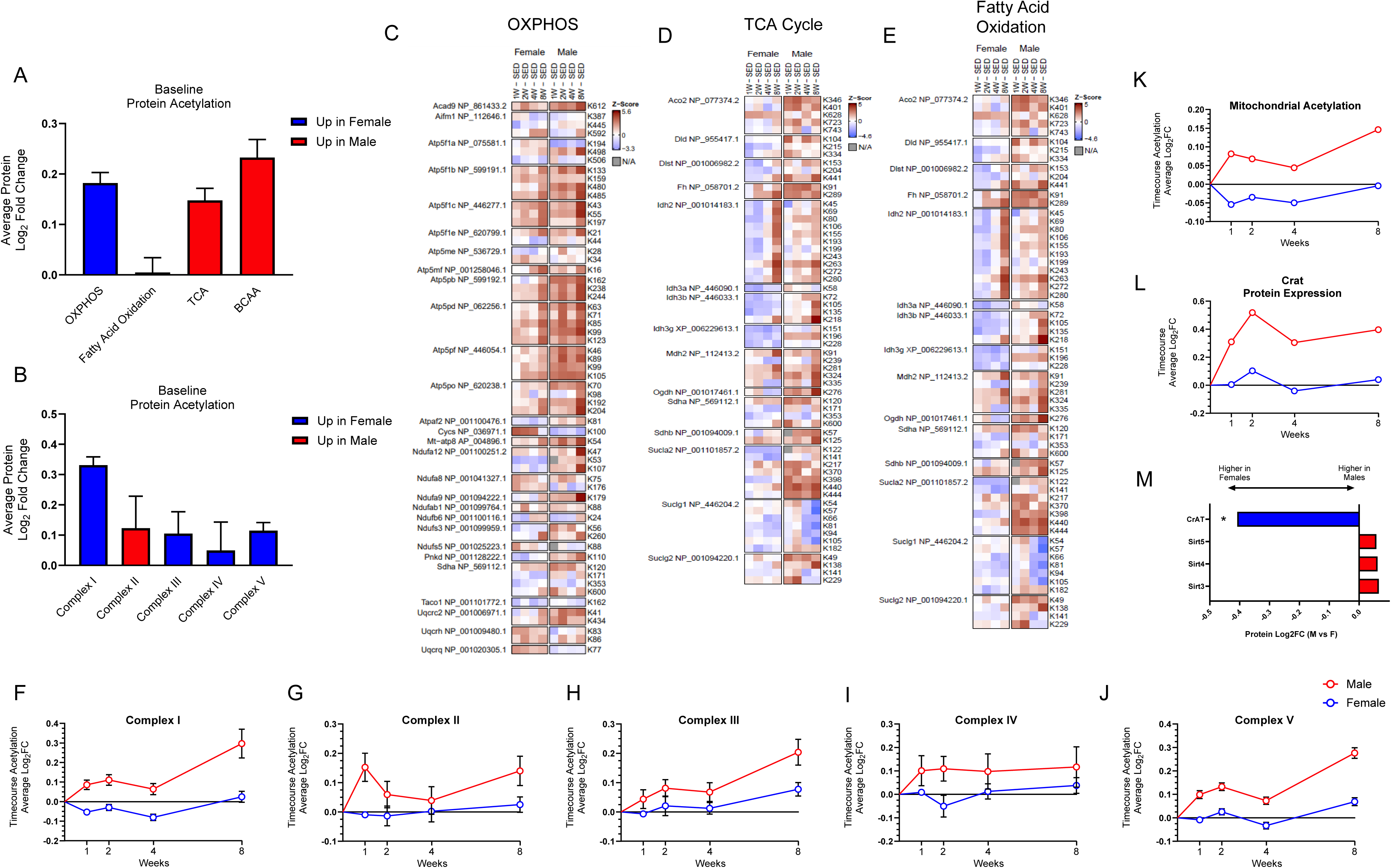
Sex-specific increase in OXPHOS, TCA cycle, and fatty acid oxidation enzymes in male liver acetylome with chronic exercise training. **A)** Average Log_2_FC of mitochondrial protein acetylation involved in OXPHOS, fatty acid oxidation, TCA Cycle (TCA), and branched chain amino acid catabolism between sedentary male and female rats. **B)** Average log_2_FC acetylation of each OXPHOS complex protein between male and female sedentary rats. **C-E)** Heat map showing the change in abundance of the most significantly altered acetylation events during each training timepoint (1wk-, 2wk-, 4wk-, 8wk-SED) in male and female rats within **(C)** OXPHOS, **(D)** TCA Cycle, and **(E)** fatty acid oxidation. **F-J)** OXPHOS complex I, complex II, complex III, complex IV, and complex V average log_2_FC in acetylation events over the course of the training paradigm. **K)** Mitochondrial acylation expressed as Log_2_FC. **L)** CrAT abundance change over the exercise training paradigm. **M)** Log_2_FC of deacetylases and acetyltransferase enzymes between sedentary male and female rats. Values are represented as mean ± standard error. For acetylation heatmaps, each contrast was filtered to the top 10 sites with the smallest p-values after filtering to the genes present in each gene set. Values are represented as mean ± standard error.

Next, we explored the specific lysine acetylation sites of OXPHOS, the TCA cycle, and fatty acid oxidation proteins. Unlike protein abundance, exercise training promoted progressive accumulation of acetylation events in male liver mitochondria (Fig. 4K), particularly on proteins involved in OXPHOS, TCA cycle, and fatty acid oxidation (Fig.4C-E, Extended Data Fig.1C-E). The influx of acetylation events on the OXPHOS system was not limited to a particular complex, with all five complexes showing increased in acetylation events through 8 weeks of exercise training, although complex I and V were particularly susceptible to increased acetylation in males (Fig. 4F-J). In contrast, acetylation levels for each of these pathways remained stable throughout 8 weeks of exercise training in female rats. While protein abundance of deacetylases Sirt3 and Sirt4 was increased following 8 weeks of exercise training period in both male and female rats^38^, male rats showed a 32% increase in the abundance of carnitine acetyltransferase (CrAT; Fig. 4L), an enzyme that catalyzes the reversible conversion of acetyl-CoA to acetyl-carnitine, to levels comparable to sedentary female rats (Fig. 4M), suggesting an innate sexual dimorphism in acetyl-CoA processing within the mitochondria.

### Liver cholesterol and bile acid (BA) biosynthesis is persistently increased in males but temporally delayed in females with chronic exercise training

Endurance exercise training analyte results (transcript and protein; FDR<0.05) were imputed into IPA to generate mechanistic networks for top-predicted biological functions and upstream regulators (absolute z-score ≥ 2; Fig.5A and B). This approach revealed exercise-induced activation of cholesterol biosynthesis as the highest enriched pathway. In addition, exercise also increased activation of OXPHOS and fatty acid beta-oxidation, among other pathways (absolute z-score ≥ 2; Fig.5A and B). IPA predicted a sex-dependent and temporal activation of the super pathway of cholesterol biosynthesis by exercise training in both the transcriptome and proteome. Specifically, exercise increased cholesterol biosynthesis across all time points in the males but was not induced by exercise in females until weeks 4 and 8 of training (absolute z-score ≥ 2; Fig. 5A and B). We further examined the liver proteome for the abundance levels of 3-hydroxy-3-methyl-glutaryl-coenzyme A reductase (HMGCR), which catalyzes a rate-limiting step for cholesterol production^69^ and sterol 27-hydroxylase (CYP27A1) the first enzyme involved in the alternative pathway of BA synthesis^70^. HMGCR and CYP27A1 were elevated in sedentary females compared to sedentary males (Fig.5C). By 8 weeks of training, CYP27A1 was elevated in the liver proteome of both males and females, with HMGCR only increased in the males (Fig.5D). No statistically significant change in liver CYP7A1 protein abundance, the rate-limiting enzyme in the classical BA synthesis pathway^71^, was observed between the sexes over 8 weeks of training; however, it tended to follow the same pattern as HMGCR abundance for both males and females (Fig.5E). Among other proteins in the cholesterol/BA synthesis pathway, CYP27A1 was also heavily acetylated by 8 weeks of training (Fig.5G). Of note, solute carrier family 27 member 5 (SLC27A5), that encodes bile acyl-CoA synthetase and aids in the conjugation of taurine and glycine to BAs^72^ and sterol carrier protein-2 (SCP-2) that stimulates liver cholesterol 7α-hydroxylase were also DAPs upregulated with exercise (Fig.2B, data not shown for SCP-2). A graphical summary of the integrated multi-omic analysis of significantly altered proteins and post-translational modifications (PTMs) for cholesterol/BA biosynthesis pathway at 8 weeks are shown in Fig.5H (females) and I (males).

**Figure 5:**
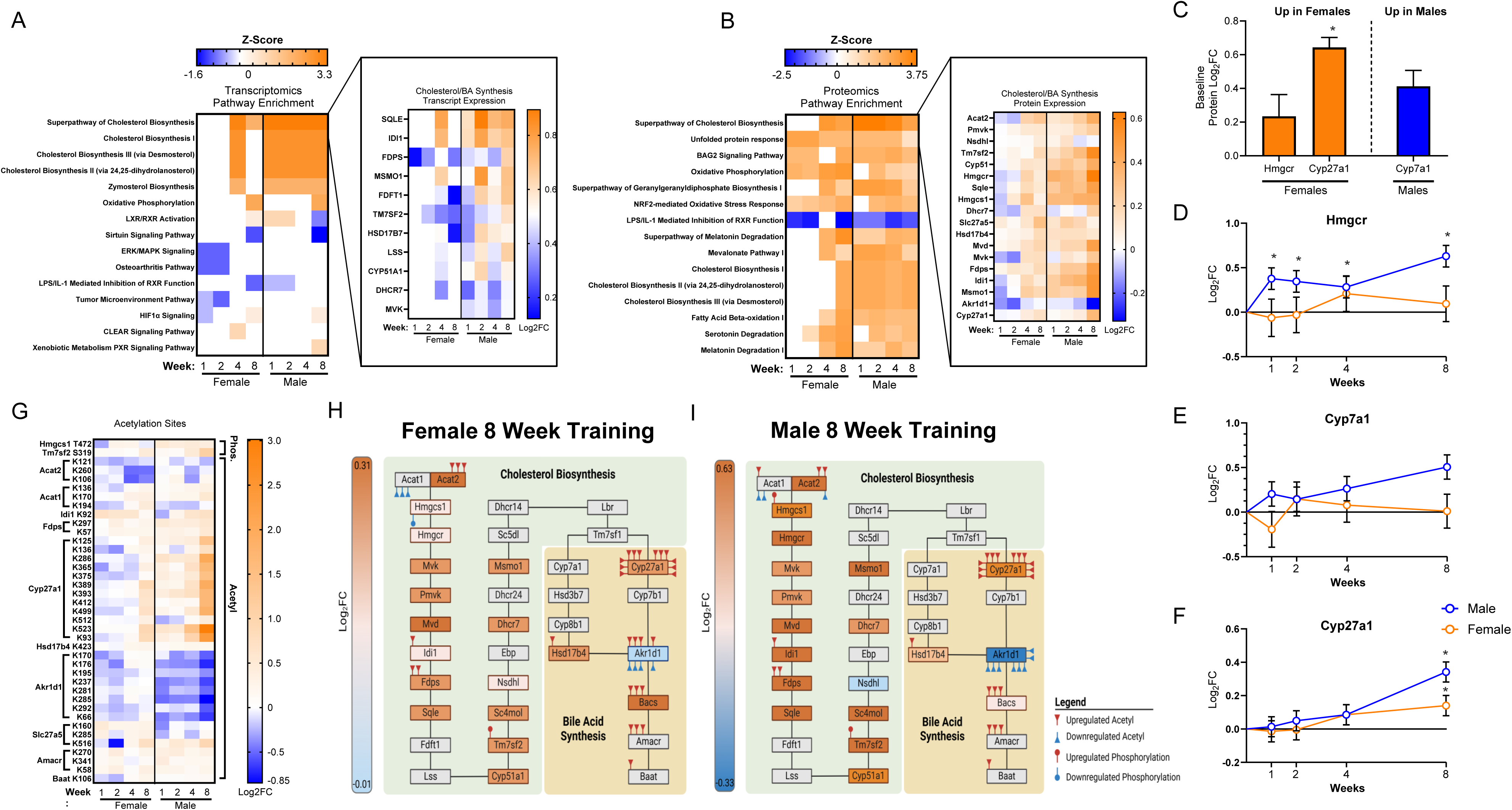
Liver cholesterol and BA biosynthesis is persistently increased in males but temporally delayed in females with chronic exercise training. **A and B)** Comparison analysis heatmaps of top regulated biological functions from IPA transcriptomic (**A)** and proteomic **(B)** analysis after 1, 2, 4, or 8 weeks of treadmill running with expression of genes associated with super pathway of cholesterol biosynthesis. **C)** Sex differences in the abundance of the major regulatory proteins of cholesterol and BA synthesis pathways in sedentary control rats. **D-F)** Timepoint specific responses to treadmill training of HMGCR **(D)**, CYP7A1 **(E)**, and CYP27A1 **(F)** at 1, 2, 4, or 8 weeks of training. **G)** Heatmap of cholesterol and BA synthesis post-translational modification (PTMs) for phosphorylation and acetylation events after 1, 2, 4, or 8 weeks of treadmill running. **H and I)** Protein and PTM visualization of cholesterol and BA synthesis pathway after 8 weeks of treadmill running in male **(H)** and female rats **(I)**. DEPs with z-score ≥ 2 were considered activated. Abundance levels of DAPs are displayed as log_2_FC. Phosphoproteome and acetylome values are expressed as log_2_FC. Blue and green colors represent a decrease and orange and red an increase in endurance training compared to sedentary rats (n=5). Values are represented as mean ± standard error. Panels H and I generated in BioRender.

Considering increases in cholesterol/BA biosynthesis were not observed in females until week 4 of training, we further explored sexual dimorphism in sedentary conditions. Sex differences in sedentary rats revealed 787 upregulated and 653 DEGs and 1512 upregulated and 1442 downregulated DAPs (FDR<0.05; Extended Data Fig. 2A and B).Utilizing IPA, compared to sedentary females, the male sedentary transcriptome predicted a decreased super pathway of cholesterol biosynthesis, cholesterol biosynthesis I, cholesterol biosynthesis III (via desmosterol), cholesterol biosynthesis II (via 24,25-dihydrolanosterol), and zymosterol biosynthesis (absolute z-score ≥ 2; Extended Data Fig.2C). Furthermore, activated upstream regulators (absolute z-score = ± 2) from transcriptomic and proteomic datasets predicted a decrease in cholesterol biosynthesis in the male sedentary rats compared with females (absolute z-score ≥ 2; Extended Data Fig.2D-E). These results show that cholesterol and BA pathways are elevated in female rats compared to males, but that both sexes experience increases with exercise training that occur on a different time course.

Further IPA analyses revealed increased activation of upstream regulators (absolute z-score ≥ 2; Extended Data Fig.3A-B) involved in synthesis of cholesterol in the liver of males and females after 8 weeks of treadmill running in females in the transcriptome. Additionally, IPA analyses revealed increased activation of upstream regulators (absolute z-score ≥ 2; Extended Data Fig.3B-D) involved in synthesis of metabolism/synthesis of phosphatidylcholine in the liver of males and females after 8 weeks of treadmill running in females.

### Chronic exercise training reduces liver lipids and changes fecal excretion of lipids in a sexual dimorphic manner

Exercise has a profound effect in lowering intrahepatic lipid stores^12,22–24,73–80^. Therefore, we also explored exercise-induced changes in pathways controlling triacylglycerol (TG) primary storage depots and intermediate pools. IPA predicted decreased concentrations of TG after 8 weeks of endurance training (Fig.6A and B). Metabolic analysis corroborated IPA transcriptomic predictions, revealing a significant decrease in liver TG, cholesterol esters (CE), phosphatidylcholines (PC), and phosphatidylethanolamines (PE) in the male livers after 8 weeks of training (p< 0.05, Fig.6C). Liver metabolomic enrichment also displayed a significant increase in C24 BAs in male livers after 8 weeks of exercise (p< 0.05, Fig.6C), with deoxycholate, hyodeoxycholate, and ursodeoxycholate composing the majority of the BA pool (Fig.6D). Deoxycholate, hyodeoxycholate, and ursodeoxycholate were quantitatively higher (not significant) across all exercise timepoints for males but peaked at week 2 for females followed by a gradual decrease. Interestingly, no significant changes in plasma lipid concentrations were detected in males. However, plasma concentration of cholesterol (One-way ANOVA, p = 0.0479), DAGs (One-way ANOVA, p = 0.001), and TGs (One-way ANOVA, p = 0.00260) were reduced with exercise in the females. Post-hoc analyses revealed cholesterol was reduced after the first week of training with DAGs and TGs reduced after 1,2,4 of training in females (p<0.05, Table 1).

**Figure 6:**
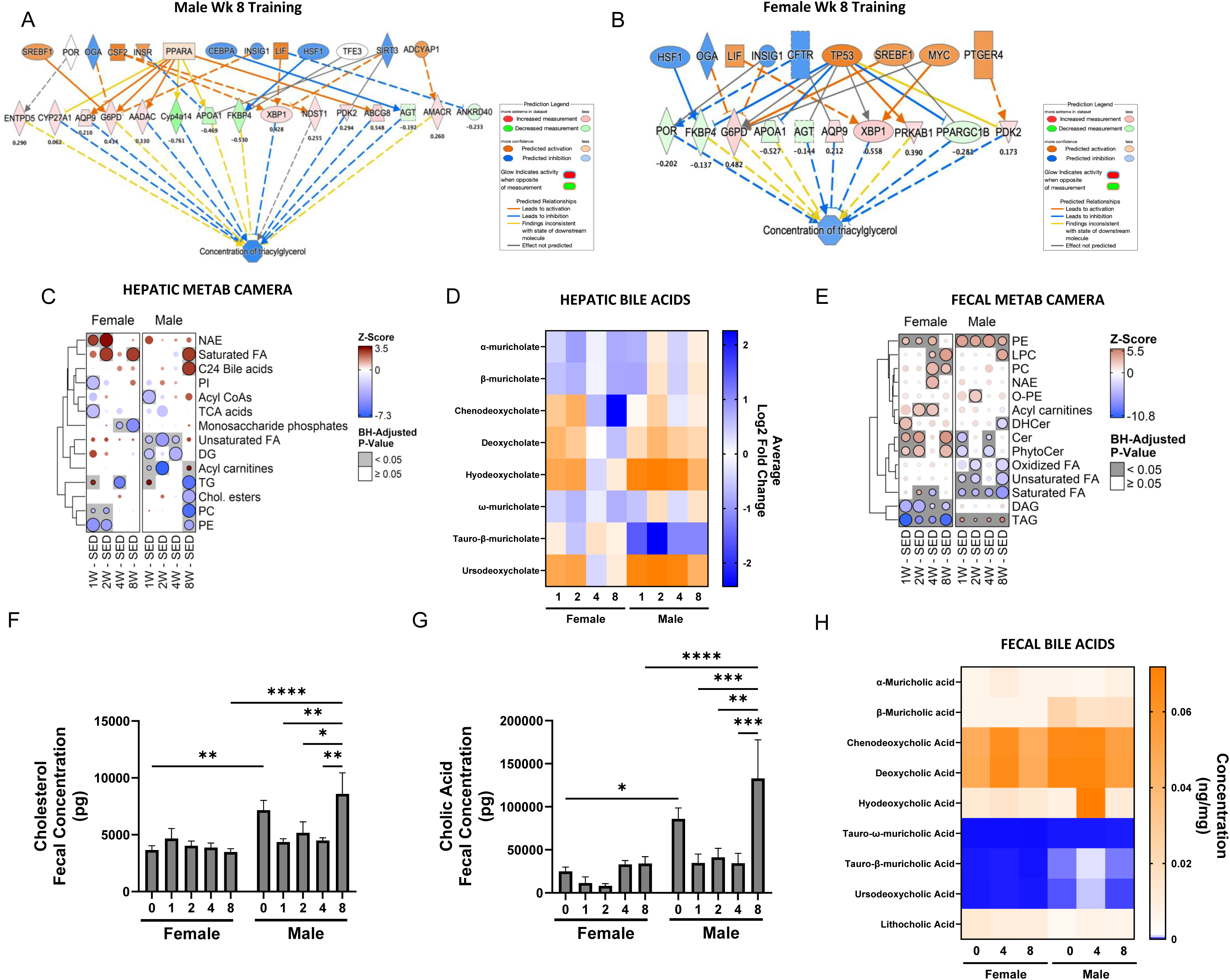
Chronic exercise training reduces lipids in the liver and changes fecal excretion of lipids in a sexual dimorphic manner. **A and B)** Transcriptomic activated upstream regulators predicted by IPA display a sex-consistent decrease in triacylglycerol concentration after 8 weeks of treadmill running in male **(A)** and female **(B)** rats. Blue and green colors represent a decrease and orange and red an increase in endurance training compared to sedentary rats (n =5). DEGs are shown below each node and displayed as log_2_ fold change. DEGs with z-score ≥ 2 were considered activated. **C)** Bubble heatmap depicting top biological functions (BP) from the Gene Ontology (GO) database across training. Heatmap was derived from the liver metabolic CAMERA-PR results methods. Circles are colored by the Z-score and scaled by row so that the most significant comparison is of maximum area. **D)** Heat map showing liver BA concentration after training (1wk-, 2wk-, 4wk-, 8wk-SED) in male and female rats (n = 5). **E)** Bubble heatmap depicting top biological processes (BP) from the Gene Ontology (GO) database across training. Heatmap was derived from the fecal metabolic CAMERA-PR results. **F-G)** Male and female fecal cholesterol **(F)** and Cholic acid **(G)** concentration at sedentary levels and after 1,2,4, and 8 weeks of treadmill running (n = 5). **J)** Fecal BA species and summated fecal BA concentration at sedentary levels and after 4 and 8 weeks of treadmill running in male and female rats (n = 7-8). A gray or black background indicates a significant result (adjusted p-value < 0.05). Values are represented as mean ± standard error.

**Table 1:**
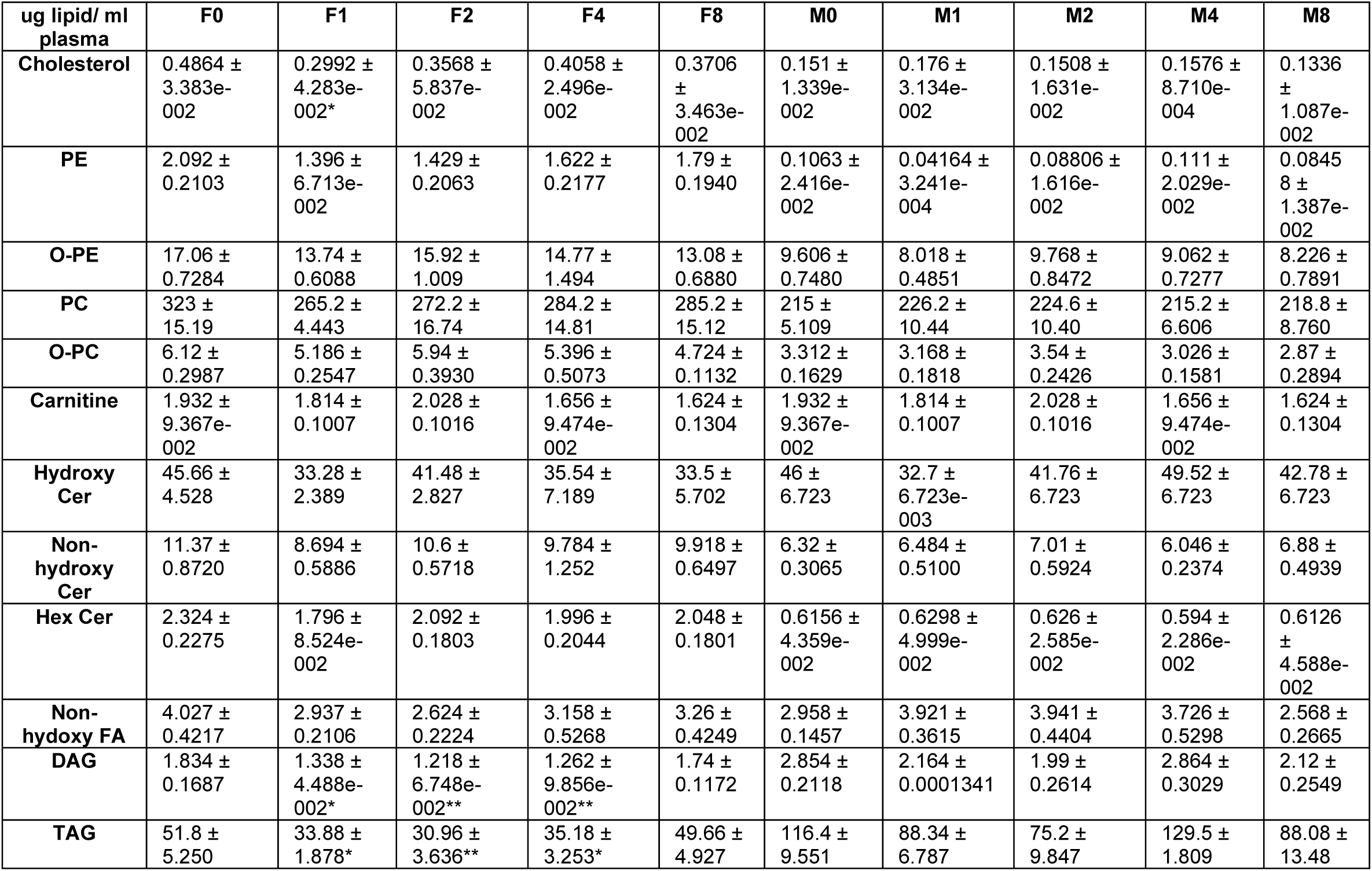
Plasma lipid concentrations in sedentary and trained male and female rats (ug/ lipid/ ml plasma). Significant protein abundance change compared to sedentary conditions within sex is presented by *, p<0.05; **, p<0.01. Values are represented as mean ± standard error.

The predicted elevation of cholesterol and BA synthesis in the liver (Fig.5) led us to investigate if an increased fecal loss of lipids, BAs, and cholesterol was causing an upregulation of synthesis to maintain systemic BA and cholesterol homeostasis. There was a significant increase in PE and lysophosphatidylcholine (LPC) with exercise (p< 0.05, Fig.6E), independent of sex. Furthermore, exercise in females also increased fecal PC, N-acylethanolamine (NAE), acyl-carnitines, and ceramides (cer) (p< 0.05, Fig.6E). There was also a sexual dimorphic response to chronic exercise. Exercise training increased fecal TGs in males but reduced them in females (p< 0.05, Fig.6E). Typically, approximately 5% of the BA pool is lost in the stool, while 95% is recycled by the liver; however, rats bred for high aerobic capacity have an approximately 50% increase in fecal BA loss compared to rats bred for low aerobic capacity^81,82^. Analyses of fecal cholesterol concentration revealed a significant interaction between sex and exercise for fecal cholesterol (two-way ANOVA, p = 0.0080) and cholic acid concentration (two-way ANOVA, p = 0.0420). Fecal cholesterol was significantly elevated after 8 weeks of exercise compared to 1, 2, and 4 weeks of exercise in males (p<0.05, Fig.6F). Moreover, fecal cholic acid concentration, a primary component of the BA pool, mirrored fecal cholesterol with a significant elevation after 8 weeks of exercise compared to 1, 2, and 4 weeks of exercise in males (p<0.05, Fig. 6G). However, females did not display increased fecal cholesterol or cholic acid. Other pools of fecal BAs were not significantly changed with exercise in either sex (Fig. 6H). These data provide convincing evidence of an exercise-induced increase in liver BA synthesis that could mediate liver and fecal cholesterol/lipid excretion in a sex specific manner.

## DISCUSSION

Over a half-century of results have clearly shown that endurance exercise significantly modulates liver metabolism^10,14^. Still, the mechanisms by which exercise exerts these effects within the liver remain largely unknown. While many studies have investigated the benefits of exercise, this is the first study to investigate the temporal, multi-omics effects of progressive exercise training within the liver in both males and females. Together, these data show that exercise training selectively increases the abundance of proteins regulating metabolic processes, particularly OXPHOS in females. In contrast, males increase the acetylation of OXPHOS, TCA cycle, and fatty acid oxidation enzymes in response to exercise training. Moreover, these data show that exercise training increases cholesterol/BA metabolism in both sexes while inducing a sexual dimorphic difference in fecal cholesterol and BA loss. Collectively, these data provide foundational clues for how exercise modulates liver metabolism and mitochondrial function in a sex-dependent manner, evidence that begins to unravel the influential role of exercise to maintain liver metabolic health and prevent or reverse MASLD.

While it has previously been reported that female rodents possess elevated liver mitochondrial protein content^83,84^ and greater individual complex protein abundance^13,34^, these analyses have relied heavily on the abundance of a single protein within OXPHOS complex proteins, or bulk quantification of proteins. This is the first study, to our knowledge, to utilize proteomics-based approaches to quantify changes in both mitochondrial protein abundance and mitochondrial acetylation events following successive weeks of endurance exercise training. At baseline, while total OXPHOS protein abundance was relatively similar, we observed significantly greater abundance of complex I proteins in sedentary female rats, which may support the expanded liver mitochondrial respiratory capacity previously reported in both female rats^83–85^ and mice compared to males^13,34^. Except complex V, female rats increased the abundance of all complexes within the OXPHOS system with exercise training to a higher degree than males. Importantly, the increase in OXPHOS protein abundance in females exceeded the average exercise-mediated increase in mitochondrial proteins, showing that OXPHOS proteins were preferentially increased in females. These changes coincided with enrichment of various other metabolic processes in female livers demonstrating that exercise training in females mediates paired increases in the rate of substrate catabolism and OXPHOS complex abundance.

Conversely, while we observed greater abundance of proteins involved in metabolic pathways such as fatty acid oxidation, branched-chain amino acids, and carbohydrate metabolism in sedentary male rats versus females, males could not sustain further exercise-meditated increases of these pathways after 2 weeks of endurance training. These findings provide mechanistic evidence for the previous findings in female mice where females show robust changes in mitochondrial respiratory capacity to metabolic stress^13^, greater mitochondrial coupling control, and an enhanced pairing of oxygen consumption to ATP production compared to males^13,32^. The differential ratio of TCA cycle and OXPHOS protein abundance between male and female rats suggests unique handling of TCA cycle intermediates. Based on innate differences in susceptibility to MASLD and liver mitochondrial respiratory capacity^13,34^, it is likely that female rats tightly couple TCA cycle flux to the OXPHOS system to stimulate energy production, while the male liver may be preferentially geared towards utilizing TCA cycle intermediates to support the synthesis of metabolites through cataplerosis, independent of paired increases in mitochondrial respiration. The coupling of TCA cycle flux to mitochondrial respiration in females may drive the observed paired increases in protein abundance involved in metabolic pathways and OXPHOS components. In males, the elevated baseline abundance of metabolic proteins suggests they may not require additional increases in protein content to match demand increases. If TCA cycle flux is indeed not coupled as tightly to mitochondrial respiration in males, this could explain the limited change in abundance of OXPHOS components. While studies investigating metabolic fluxes in the liver following exercise are limited, data in prolonged high-fat diet fed male rats supports the notion that males are capable of increased TCA cycle cataplerosis and anaplerosis independent of elevations in mitochondrial respiration^87^.

However, males demonstrably showed a robust increase in acetylation of mitochondrial proteins following exercise training. In the absence of changes in protein abundance for OXPHOS complex proteins, increased substrate load within the mitochondria both during and post-exercise training may promote the accumulation of substrates, including acetyl-CoA, which the mitochondria must find ways to mitigate. In the present dataset, there was a robust elevation of CrAT in male rats across all training timepoints that was not observed in the females. Classically, CrAT mediates export of acetyl groups from the mitochondrial matrix by converting acetyl-CoA to acetyl carnitine, regulating the levels of free CoA pools. However, evidence in skeletal muscle suggests that exercise can reverse normal CrAT-mediated acetyl carnitine flux, preferentially promoting increased acetyl-CoA levels^87^. Due to the bidirectional activity of CrAT, we postulate that CrAT acetylation in males may serve as a mechanism by which male rats preferentially accumulate acetyl-groups within the mitochondria, exceeding the capacity of mitochondrial deacetylases^38^ and increasing mitochondrial acetylation via mass action. Ketone body synthesis is another pathway linked to removing excess acetyl-CoA from the mitochondria, particularly during exercise. However, we observed a significant downregulation of protein abundance through the ketone body synthesis pathway, further supporting our hypothesis that acetylation likely occurred due to a downregulation of pathways that aid in mitochondrial matrix acetyl CoA export^88^.

In addition to these mitochondrial effects, endurance exercise training displayed a robust increase in analytes associated with liver cholesterol biosynthesis across multiple omes that were more robust in males than females. The rate-limiting enzyme for cholesterol biosynthesis, HMGCR^69^, was upregulated in males at all exercise timepoints compared to sedentary conditions, supporting exercise-induced elevation in cholesterol biosynthesis. Differentially expressed genes/proteins of cholesterol biosynthesis (including HMGCR) were lower in sedentary males compared to sedentary females, a finding consistent with previous reports^89^. We posit that for any complex trait elevated in a sex-biased manner under sedentary conditions (e.g., cholesterol biosynthesis), a higher volume or duration of exercise may be required to induce an adaptation.

Several lines of evidence in this study support that endurance exercise significantly mediates liver cholesterol and BA synthesis. First, liver transcriptomic enrichment analysis using CAMERA revealed sterol-related biosynthetic processes were upregulated across all exercise timepoints. Second, transcriptomic and proteomic enrichment analysis using IPA revealed the super pathway of cholesterol and BA synthesis as top canonical pathways responsive to exercise training. Moreover, investigation in the liver proteome found the rate-limiting enzyme for alternative BA synthesis: CYP27A1,^70^ was of higher abundance and was highly acetylated in both males and females after 8 weeks of exercise. Exercise training also elevated protein levels of SLC27A5, which encodes bile acyl-CoA synthetase and aids in the conjugation of taurine and glycine to BAs^72^, and carrier protein-2 (SCP-2), which regulates CYP7a1 (the rate-limiting enzyme in liver BA synthesis^90^). In addition, liver metabolomic analysis displayed a robust increase in liver BA synthesis after 8 weeks of training in male rats. Together, these data provide compelling evidence for exercise-mediated increases in liver cholesterol and BA synthesis in male rats with moderate effects in females that required a longer duration of exercise training to occur.

Cholesterol is a substrate for BA synthesis and fecal loss of BAs (approximately 5% per day) is a primary method of cholesterol elimination^91^. Abundance profiles suggestive of increased liver BA synthesis were elevated after 8 weeks of training in male rats, coinciding with reductions in several liver lipid species: TGs, CE, PE, and PC. Furthermore, TG, PE, and total cholesterol were elevated in the fecal along with increased cholic acid, a primary BA that plays a significant role in excretion of cholesterol from the body^91^. Furthermore, 8 weeks of exercise decreased male liver TG concentration, a well-known adaptation with exercise^79^, with minimal effects on plasma lipid concentrations. We hypothesize this strong association between cholesterol and cholic acid could be a mechanism for exercise-induced reductions in liver cholesterol in males. These data corroborate previous findings that exercise training (combined with diet-induced weight loss) increased markers of liver BA synthesis^92^ and that high aerobic capacity and exercise are associated with upregulated BA synthesis and greater fecal excretion of cholesterol/BA in rats fed a high fat diet paired with protection against steatossi^81^. These links are further supported by evidence that BA sequestrants (mediate increased fecal BA loss) reduce liver lipids in mice^93,94^, suggesting exercise-mediated fecal BA loss may have a significant role in the treatment of steatosis.

Decreased proliferation of hepatic stellate cells/fibrosis was identified as a cell-type specific adaptation with exercise. Many DEGs identified were found in both male and female liver transcriptomes, suggesting chronic exercise decreased hepatic stellate cell activation, independent of sex. The genes identified in both males and females included AGT and CCND1. AGT is converted to angiotensin in the liver^95^, which promotes HSC activation and fibrosis through profibrotic pathways such as TGF-β1 signaling and oxidative stress^96,97^. Overactivation of the renin-angiotensin system (RAS) is a known driver of fibrogenesis that exercise protects against^98^, potentially through downregulation of AGT. CCND1 regulates cycle progression from G1 to S phase and is associated with proliferation and activation of HSCs^99^. Exercise can revitalize quiescent skeletal muscle stem cells through Cyclin D1^100^, but this is the first study to report chronic exercise decreases CCND1 protein abundance in the liver. Considering that single cell sequencing was not performed, changes could be occurring in cell types other than HSCs in the liver. These data provide potential genes that modulate exercise and decreased stellate cell activation.

These studies provided valuable insight into liver adaptations in response to exercise across 1,2,4,8 weeks of exercise that would be unattainable in humans participants. Although a rat model was utilized in these studies despites aligning with several of the observations from humans [25677548], the use of rodents as a translational model presents inherent challenges in directly extrapolating findings to human physiology [33180643]. Differences in metabolic rate[28367138], muscle fiber composition[22530000] between rats and humans may influence the observed adaptations to exercise. Additionally, while these studies comprehensively assessed molecular markers of exercise adaptation, this study did not include functional measurements such as mitochondrial respiration or metabolic flux that could have been correlated against molecular outcomes. Furthermore, these studies provided data 48 hours post training, missing the acute response to exercise and did not consider other exercise modalities that could be addressed in future studies[32589957].

The combination of multi-omic approaches (transcriptome, proteome, phosphoproteome, metabolome, and lipidome), time course, and inclusion of both biological sexes in this data set provides an unprecedented perspective on the liver adaptations to endurance exercise training. Moreover, the results point to a marked sexual dimorphism in liver metabolism and mitochondrial profiles both in sedentary conditions and in temporal responses to endurance exercise training. Several exercise-induced adaptations only occur after longer durations of training, suggesting that for hepatoprotective effects, long-term endurance is likely most efficacious. The multi-omic data mining tools found at the bioinformatics core (MoTrPAC BIC; https://motrpac-data.org) can be leveraged for hypothesis-driven mechanistic studies to understand further the effects of exercise training on the liver, particularly at the mitochondrial level. The robust exercise-induced changes in OXPHOS, cholesterol/BA synthesis, and mitochondrial acetylation point to novel targets which can be leveraged to combat metabolic diseases centered in the liver, including MASLD.

## METHODS

### Animals

Adult male and female Fischer 344 inbred rats were acquired from the National Institute of Aging (NIA) rodent colony. Upon arrival, rats were adapted to a 12-hr reverse light-cycle, ensuring treadmill training occurred during the active period. Rats were housed as same-sex pairs in ventilated racks (Thoren Maxi-Miser IVC Caging System) on Tekland 7093 shredded Aspen bedding and fed ad libitum with Charles River Rat and Mouse 18% (Auto) 5L79 LabDiet (Gateway Lab Supply, St. Louis, Missouri), which has the following calorie composition: 21.196% protein, 14.774% fat (ether extract), 64.030% carbohydrates. These are the standard bedding and diet used at the NIA rodent colony. The animal housing room was monitored daily and was maintained at a temperature of °F and relative humidity of 25–55%. Red lights were used during the dark cycle to provide adequate lighting for routine housing tasks, rodent handling, and training. All animal procedures were approved by the Institutional Animal Care and Use Committee at the University of Iowa, where the rats were housed. Complete description of animal husbandry described previously^37^.

### Treadmill familiarization and training

Treadmill familiarization and training was conducted on a Panlab 5-lane rat treadmill (Model LE8710RTS, Harvard Instruments). All animal handling and exercise sessions were performed in the active, dark phase of the rats. Following acclimation, rats underwent familiarization for 12 days to acclimate rats to the treadmill and identify potential non-compliant rats. Rats that were unable to run on the treadmill for 5 minutes at a speed of 10 m/min and grade of 0° were classified as non-compliant and removed from the study. Rats that successfully completed the 12-day familiarization protocol were randomly assigned to a training or control group. Training began at 6 months of age and lasted 1,2,4, or 8 weeks. Rats were exercised for 5 consecutive days per week using a progressive protocol designed to maintain an intensity of 70% of VO_2_ max (increasing grade and speed, see MoTrPAC^35^ for details) reaching 50min/day at a 10° grade. To control for any non-exercise-related treadmill effects, sedentary control rats were placed on the treadmills for 15 min/day at a speed of 0 m/min for 5 consecutive days per week, following a schedule similar to the 8-week-trained rats. Rats unable to exercise for at least 4 days per week were removed from the study.

### Tissue collection

Tissues were collected 48 hours following the last exercise bout. On the day of collection, food was removed at 8:30 am, 3 hours prior to the start of dissections, which occurred between 11:30 am and 2:30 pm (in the dark cycle). Rats were anesthetized with 1-2% isoflurane, blood was drawn via cardiac puncture, the liver and fecal (from the rectum) were removed and immediately flash-frozen in liquid nitrogen and stored at −80°C. Cardiac exsanguination resulted in death. Rat tissues were archived at the MoTrPAC Biospecimens Repository, until distributed to Chemical Analysis Sites for respective assays. For a more detailed description please see associated MoTrPAC publication^35^.

### Multi-omic data generation and processing

For complete detailed descriptions of methods used for sample preparation and multi-omics data generation and processing at chemical analysis sites for this study (transcriptomics, proteomics, phosphoproteomics and metabolomics/lipidomics), please see the associated MoTrPAC publication^35^. Sample preparation and multi-omics data generation and processing were performed the same as the associated MoTrPAC publication^35^ except where specified. At all multi-omic analysis sites, an unblinded batching officer managed the randomization of samples across appropriately sized batches for available analysis platforms. These randomized samples were blinded to all individuals involved in sample preparation, data generation, and initial data processing. Downstream quality control and data analysis were conducted with knowledge of experimental conditions.

### Untargeted Lipidomics

Untargeted lipidomic analysis were performed as previously described^35^. We performed semi-absolute quantification of the annotated lipids. The concentrations of each lipid species were calculated by normalizing their peak area by that of the internal standard (ISTD) from its own class and the same ESI mode and then multiplying by the known ISTD concentration. The closest match was selected when a ISTD was not detected or if no ISTD available. After calculating the relative lipid abundance, we calculated the lipid subclass sums for each sample and identified outliers on the lipid subclass level within each ESI mode. Sample-level lipid sub-classes that were more than 5 median absolute deviations from the median were identified as outliers, and the entire lipid class was removed. The missing values were imputed using the NIPALS algorithm from the mixOmics package^42^. The resulting negative imputed values for 19 TG species were replaced by feature mean. ISTD-normalized data are reported as μg/mg tissue for liver, μg/mg for plasma, or pg for fecal lipidomics.

### Fecal Lipidomics

Fecal samples (100 mg) were first homogenized with 1 mL homogenizer solution consisting of a 1:4 volumetric ratio of water:methanol. Samples were then centrifuged at 10000g for 5 minutes at 4 °C. The supernatant was then plated. Using the Biotage Extrahera, a methanol crash was performed with Solid Phase Extraction. 1500 μL of Methanol was added to the samples prior to eluting the samples through a methanol preconditioned filter. The collected eluant was then dried with nitrogen gas and constituted in 200 μL of 1:1 volumetric ratio of Acetonitrile:Methanol and stored at −80 °C until analysis. Chromatography was run on Agilent 6495C with 1290 Infinity II on Agilent Zorbax Eclipse Plus C18 (2.1 x 100 mm, 1.8 µm) column at flow rate of 0.5 mL/min at 65 °C during a 21 minute gradient. The mobile phase of UPLC grade solvents consisted of solvent A: 0.1% formic acid in water and Solvent B: 0.1% formic acid in acetonitrile. The injection volume of sample was 5 µL. Analysis was completed in negative mode. The Nozzle Voltage at −2000 V, Sheath Gas Flow at 11 L/min and Sheath Gas Temp at 400 °C. Elution gradients and transition list are provided below. Sample data was calibrated against an external standard solution that was used to optimize instrumental parameters and establish limits of detection. Peak determination, peak area integration, and calculation of calibration curves for standards was performed with the Agilent MassHunter Workstation version 10.1.

### Differential analysis

Differential analysis of the multi-omics datasets was performed as described by MoTrPAC^35^. Since sample variability appeared to be influenced by sex, with higher variability in female samples, male and female datasets were analyzed separately. Each training time point was compared against their sex-matched sedentary controls, and males were compared to females at the sedentary and 8-week-trained timepoints.

For RNA sequencing, filtered raw counts were input in accordance with DESeq2 workflow to obtain DEG’s^43^. For proteomics, limma with empirical Bayes variance shrinkage was used to obtain DAPs^44^. Specifically, likelihood ratio tests (DESeq2::nbinomLRT, lrtest) or F-tests (limma) were used to compare the model with ome-specific technical covariates and training groups as a predictor variable (i.e., sedentary control, 1 week, 2 weeks, 4 weeks, 8 weeks) against a reduced model with only technical covariates. All metabolomics datasets were log_2_-transformed and analytes with > 20% missing values were removed. Median sample–sample correlation was used to identify outlier samples, which were manually reviewed by the metabolomics sites. Due to the overlap in the coverage of different metabolomics and lipidomics assays, some metabolites/lipids were measured in multiple platforms. Redundant metabolites/lipids were curated by taking into consideration their properties (polarity, solubility, etc.) and their respective assay methodologies (extraction solvent, elution solvent, column, etc.). Specific curation steps for overlapping coverage are described in detail elsewhere^45^.

Male- and female-specific p-values were combined using Fisher’s sum of logs meta-analysis to generate differentially expressed analytes. These p-values were adjusted across all datasets within each ome to control the FDR using independent hypothesis weighting as in previous MoTrPAC publications^35^. Training-differential features were selected at 5% FDR. Covariates were selected from assay-specific technical metrics that explained variance in the data and were not correlated with exercise training (i.e., RNA integrity number (RIN), median 5’-3’ bias, percent of reads mapping to globin, and percent of PCR duplicates as quantified with Unique Molecular Identifiers).

### Correlation Adjusted MEan RAnk gene set testing

The differential analysis results were collapsed to the gene (proteomics and transcriptomics), single phosphorylation site (phosphoproteomics), or Reference Set of Metabolite Names (RefMet) level by selecting the most extreme z-score for each combination of target feature and contrast. If any source features did not map to a target feature^46^, the source feature ID was used to avoid unnecessarily removing data. These z-scores, along with Gene Ontology Biological Process gene sets^47,48^, from the Molecular Signatures Database (MSigDB)^49,50^, phosphorylation sites grouped according to their known kinases from PhosphoSitePlus^51^, and RefMet IDs grouped according to their RefMet chemical subclasses were used as input for CAMERA-PR. CAMERA-PR is a modification of the two-sample t-test that accounts for inter-molecular correlation to more correctly control the false positive rate^52^. CAMERA-PR was performed with the cameraPR.matrix function from the TMSig R/Bioconductor package (10.18129/B9.bioc.TMSig)^53,54^.

### Mitochondrial Proteome Identification

MitoCarta3.0 was referenced to isolate all mitochondrial proteins and the pathways they associate with from the entire liver proteome identified by the untargeted and acetyl-proteomics analysis, as previously described^30,55^. To determine overall directional abundance of a given pathway or protein complex, the log_2_ fold-changes of all proteins within a pathway were averaged. This analysis was performed across-sex at sedentary levels, and within-sex for each training timepoint.

### Ingenuity pathway analysis enrichment

Additional pathway and network analysis were performed using IPA software (QIAGEN Inc., Redwood City, California), similar to previous studies^56–61^. Differentially expressed analytes (i.e genes or proteins) and corresponding log_2_ fold changes were used as IPA input. IPA compared uploaded data to a curated knowledge base of biological interactions, pathways, and molecular relationships derived from peer-reviewed literature to identify enriched pathways and upstream regulators based on overrepresented genes/protein clusters in the dataset. IPA was used to generate interactive visualizations (pathway diagrams, network, and heatmaps) and determines statistical significance based on p-value and z-scores of these pathways and upstream regulators. The canonical pathway tool was used to identify overrepresented pathways with a -log (p-value) threshold of 1.3 (Fischer’s Exact Test) and an absolute z-score > 2 was used to determine pathway enrichment. Upstream Regulator analysis tool predicted transcription factors, kinases, and other regulatory molecules responsible for observed gene and protein abundances changes. Comparison analysis function in IPA was utilized to compare datasets from each timepoint and sex comparison to identify significant (absolute z-score>2) biological pathways. The sex-matched 8-week training differential features were further analyzed by identification of upstream regulators that were used to build disease and function pathways for the proteomic dataset. Canonical pathways and upstream regulators with a z-score exceeding ± 2 generated by IPA’s proprietary prediction algorithm were considered significant (i.e, activated or inhibited)^62^.

### Statistical analysis software

The R programming language (A Language and Environment for Statistical Computing. R Foundation for Statistical Computing, Vienna, Austria. https://www.R-project.org) was used to perform statistical analyses and generate most figures with the remainder created with IPA (Qiagen Inc., Redwood City, California USA, https://digitalinsights.qiagen.com) or GraphPad Prism (GraphPad Software, Boston, Massachusetts USA, www.graphpad.com). Data and analysis tools for the MoTrPAC landscape paper^35^ are also provided through the MotracRatTraining6moData and MotrpacRatTraining6mo R packages, respectively (github.com/MoTrPAC/MotrpacRatTraining6moData, github.com/MoTrPAC/MotrpacRatTraining6mo); the former package was used to access data for MotrpacRatTraining6moWATData (github.com/PNNL-Comp-Mass-Spec/MotrpacRatTraining6moWATData). Statistical analyses for fecal cholesterol and cholic acid concentration were performed with GraphPad Prism version 10.1.2 (Prism) with α=0.05. Two-way analysis of variance (ANOVA) was used with a Tukey test for multiple comparison procedure. One-way ANOVA was used for plasma lipids compared to sedentary (F0 or M0) state, respectively, with a Dunnett’s test for multiple comparison procedure. Bubble heatmaps were generated with the enrichmap function from the TMSig (10.18129/B9.bioc.TMSig) R/Bioconductor package^53^. BioRender was used to generate Fig. 3I and Fig.5. H-I.

## Data Availability

MoTrPAC data is publicly available at motrpac-data.org/data-access. Data access inquiries should be sent to. Additional resources can be found at motrpac.org and motrpac-data.org.

## Code Availability

MoTrPAC data processing pipelines for RNA-Seq and proteomics is available at: https://github.com/MoTrPAC/motrpac-rna-seq-pipeline and https://github.com/MoTrPAC/motrpac-proteomics-pipeline. Normalization and QC scripts will be available at https://github.com/MoTrPAC/motrpac-bic-norm-qc. Code for the underlying differential analysis for the manuscript will be provided in the MotrpacRatTraining6mo R package (motrpac.github.io/MotrpacRatTraining6mo).

---

**Extended Data Table 1:** List of abbreviations and their meanings.

|  |  |
| --- | --- |
| MASLD | Metabolic-Dysfunction-Associated Steatotic Liver Disease |
| HMGCR | 3-Hydroxy-3-Methyl-Glutaryl-Coenzyme A Reductase |
| ANOVA | Analysis Of Variance |
| AGT | Angiotensinogen |
| BA | Bile Acid |
| BP | Biological Processes |
| Bckdha | Branched Chain Keto Acid Dehydrogenase E1 Subunit Alpha |
| Bckhdb | Branched Chain Keto Acid Dehydrogenase E1 Subunit Beta |
| CrAT | Carnitine Acetyltransferase |
| CDC7 | Cell Division Cycle 7 |
| Cer | Ceramides |
| CE | Cholesterol Esters |
| CCND1 | Cyclin D1 |
| DEGs | Differentially Expressed Genes |
| DAPs | Differentially Abundant Proteins |
| FDR | False Discovery Rate |
| GO | Gene Ontology |
| HSC | Hepatic Stellate Cell |
| Hadha | Hydroxyacyl-CoA Dehydrogenase Trifunctional Multienzyme Complex Subunit Alpha |
| Hadhb | Hydroxyacyl-CoA Dehydrogenase Trifunctional Multienzyme Complex Subunit Beta |
| IPA | Ingenuity Pathway Analysis |
| ISTD | Internal Standard |
| KSEA | Kinase–Substrate Enrichment Analysis |
| LPC | Lysophosphatidylcholine |
| mTOR | Mammalian Target of Rapamycin |
| MAP | Mitogen-Activated Protein |
| MSigDB | Molecular Signatures Database |
| MoTrPAC | Molecular Transducers of Physical Activity Consortium |
| NAE | N-Acylethanolamine |
| NIA | National Institute of Aging |
| OXPHOS | Oxidative Phosphorylation |
| PC | Phosphatidylcholines |
| PE | Phosphatidylethanolamines |
| PTMs | Post-Translational Modifications |
| CAMERA-PR | Pre-Ranked Correlation Adjusted Mean Rank Gene Set Testing |
| Pdha1 | Pyruvate Dehydrogenase E1 Subunit Alpha 1 |
| Pdhb | Pyruvate Dehydrogenase E1 Subunit Beta |
| RefMet | Reference Set Of Metabolite Names |
| RAS | Renin-Angiotensin System |
| RIN | RNA Integrity Number |
| SLC27A5 | Solute Carrier Family 27 Member 5 |
| CYP27A1 | Sterol 27-Hydroxylase |
| SCP-2 | Sterol Carrier Protein-2 |
| SREBP1c | Sterol Regulatory Element Binding Protein 1c |
| Sulcg1 | Succinate-CoA Ligase Gdp/Adp-Forming Subunit Alpha |
| Sulcg2 | Succinate-CoA Ligase Gdp/Adp-Forming Subunit Beta |
| TG | Triacylglycerol |
| TGF-β1 | Transforming Growth Factor Beta-1 |

**Extended Data Figure 1:**
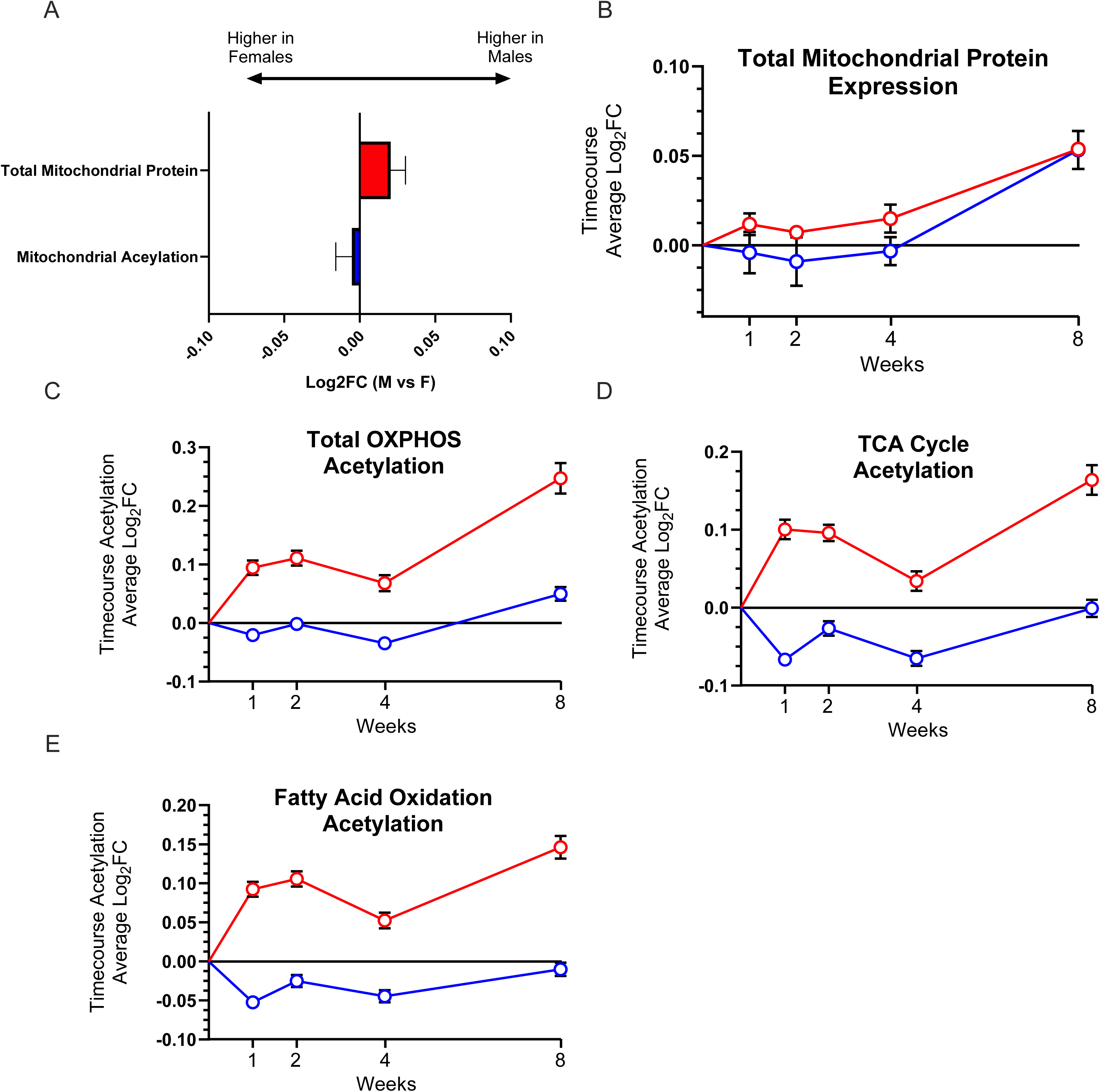
Sedentary and exercise-mediated changes in the abundance of mitochondrial proteome and acetylome in the liver. **A)** Average log_2_FC of total mitochondrial protein abundance and acetylation events between sedentary male and female rats. **B)** Exercise-mediated change in mitochondrial protein abundance presented as average log_2_FC for all mitochondrial protein. **C-E)** Pathway specific average log_2_FC in protein acetylation for OXPHOS, TCA cycle and fatty acid oxidation. **G)** CrAT abundance change over the exercise training paradigm. **H)** Log_2_FC of deacetylases and acetyltransferase enzymes between sedentary male and female rats. Values are represented as mean ± standard error.

**Extended Data Figure 2:**
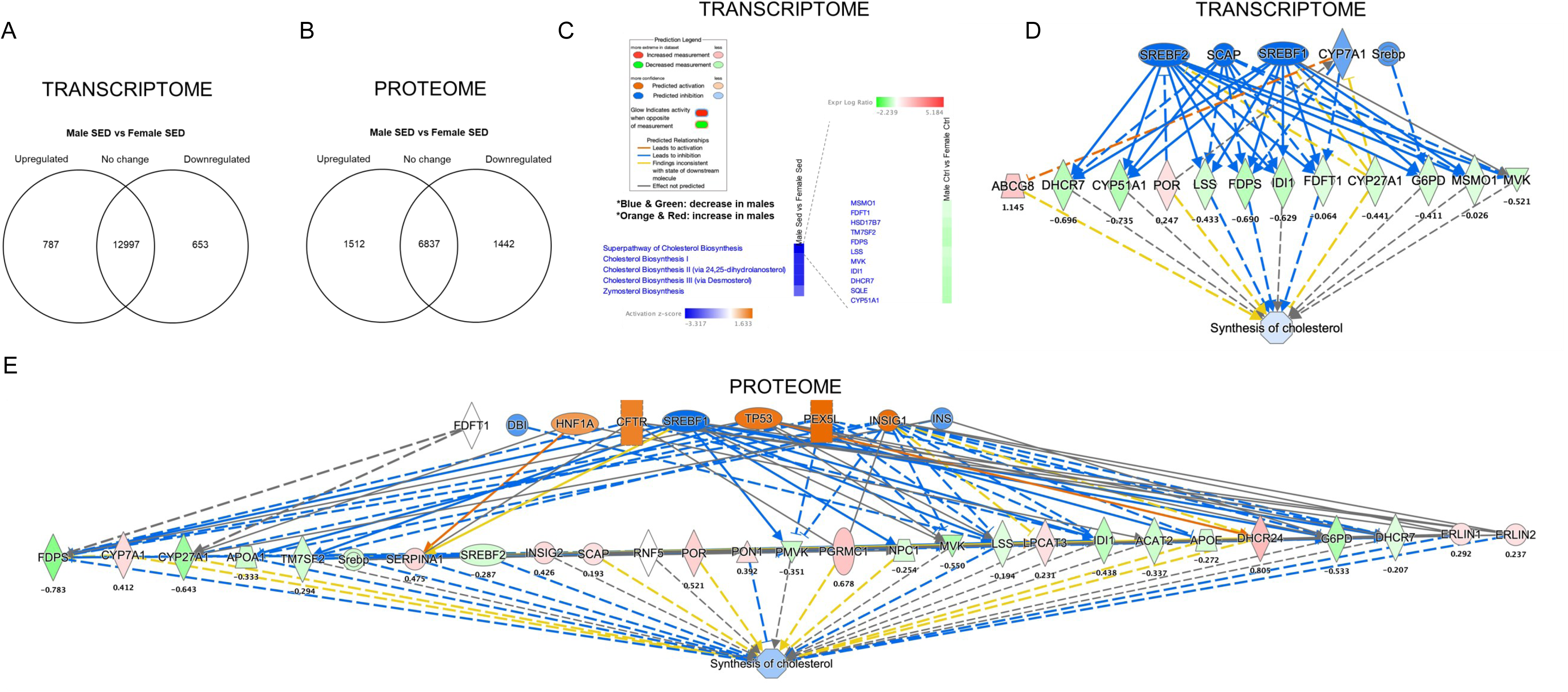
Transcriptome and proteome predicted reduced cholesterol biosynthesis in the liver of male compared to female sedentary rats. **A and B)** Venn diagram of differentially expressed genes (DEGs) and differentially abundant proteins (DAPs) in male verse female sedentary rats. **C)** Comparison analysis heatmaps of top regulated biological functions from IPA transcriptomic analysis in male verse female sedentary rats (bottom left panel) and genes associated with super pathway of cholesterol biosynthesis (right panel). IPA figure legend (upper left panel). **D and E)** Transcriptomic and proteomic activated upstream regulators predicted by IPA display downregulated cholesterol synthesis in male versus female sedentary rats. DEGs and DAPs take all timepoints and sexes into account (FDR<0.05). Blue and green colors represent a decrease and orange and red colors represent an increase in males compared to female sedentary rats. DEGs and DAPs are shown below each node and displayed as Log_2_FC. For DEGs and DAPs, a z-score ≥ 2 was considered activated.

**Extended Data Figure 3:**
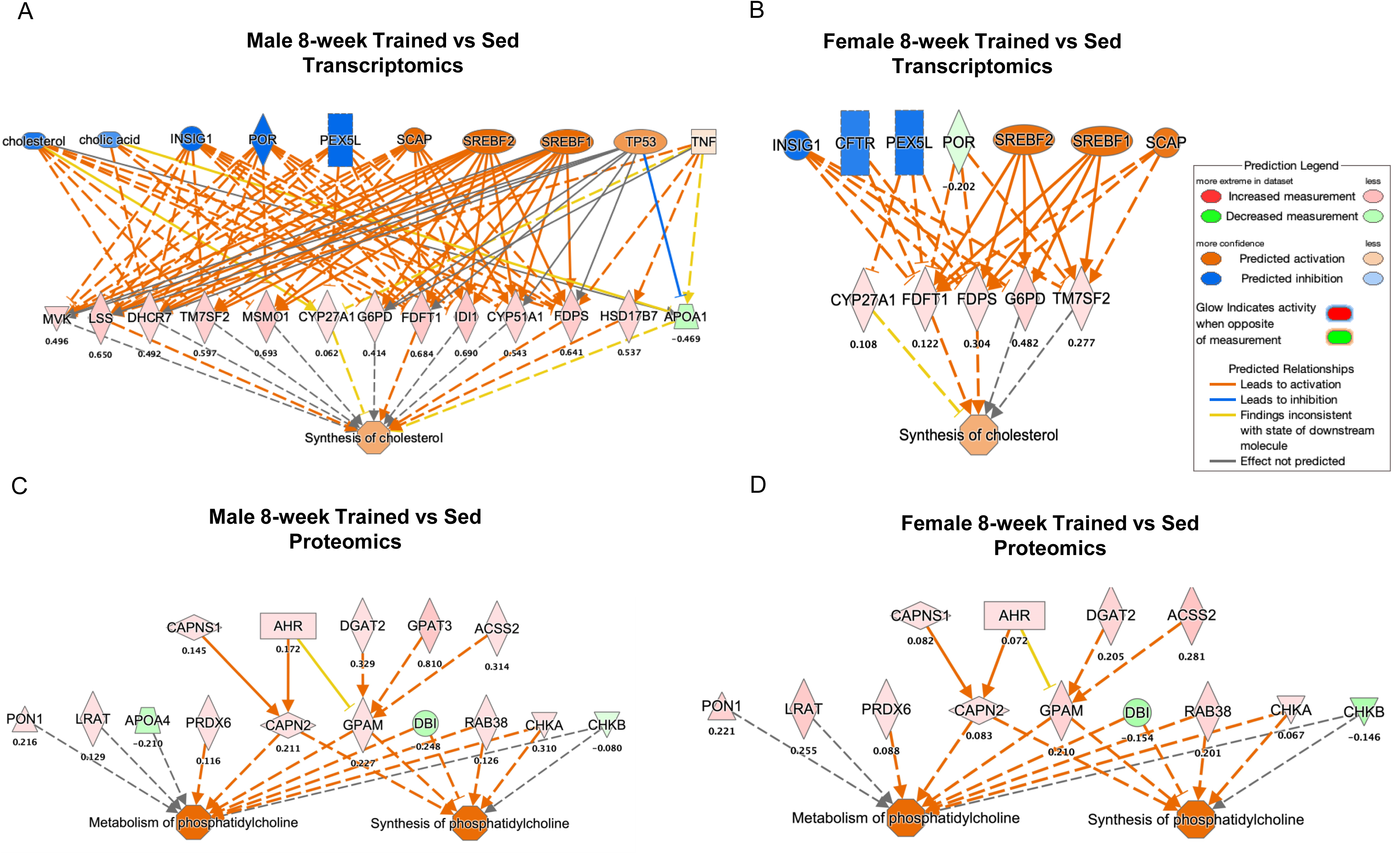
Transcriptome and proteome predicted enhanced lipid metabolism after 8 weeks of exercise in the liver, independent of sex. **A and B)** Predicted increased synthesis of cholesterol in the liver of **(A)** males and **(B)** females after 8 weeks of treadmill running. **C and D)** Predicted increased synthesis of metabolism/synthesis of phosphatidylcholine in the liver of **(A)** males and **(B)** females after 8 weeks of treadmill running. DEGs take all timepoints and sexes into account (FDR<0.05). Blue and green colors represent a decrease and orange, and red colors represent an increase in endurance training compared to sedentary rats (n=5). Expression levels of DEGs are shown below each node and displayed as log_2_FC (fold change). DEGs with z-score ≥ 2 were considered activated.

